# Residue-based correlation between equilibrium and rate constants is an experimental formulation of the consistency principle for smooth structural changes of proteins

**DOI:** 10.1101/2022.10.24.513469

**Authors:** Daisuke Kohda, Seiichiro Hayashi, Daisuke Fujinami

## Abstract

The consistency principle represents a physicochemical condition requisite for ideal protein folding. It assumes that any pair of amino acid residues in partially folded structures has an attractive short-range interaction *only if* the two residues are in contact within the native structure. The residue-specific equilibrium constant, *K*, and the residue-specific rate constant, *k* (forward and backward), can be determined by NMR and hydrogen-deuterium exchange studies. Linear free energy relationships (LFER) in the rate-equilibrium free energy relationship (REFER) plots (i.e., log *k* vs. log *K*) are widely seen in protein-related phenomena, but our REFER plot differs from them in that the data points are derived from one polypeptide chain under a single condition. Here, we examined the theoretical basis of the residue-based LFER. First, we derived a basic equation, ρ_*ij*_ = ½(ϕ_*i*_ + ϕ_*j*_), from the consistency principle, where ρ_*ij*_ is the slope of the line segment that connects residues *i* and *j* in the REFER plot, and ϕ_*i*_ and ϕ_*j*_ are the local fractions of the native state in the transient state ensemble (TSE). Next, we showed that the general solution is the alignment of the (log *K*, log *k*) data points on a parabolic curve in the REFER plot. Importantly, unlike LFER, the quadratic free energy relationship (QFER) is compatible with the heterogenous formation of local structures in the TSE. Residue-based LFER/QFER provides a unique insight into the TSE: A foldable polypeptide chain consists of several folding units, which are *consistently* coupled to undergo smooth structural changes.

Significance
The physicochemical basis of smooth protein folding has been theoretically explained by the consistency principle. We propose that the consistency principle is formulated by the quadratic relationship in the double logarithm plot of the residue-specific equilibrium and rate constants of a polypeptide chain. The quadratic relationship offers a procedure for the experimental verification of the consistency principle. One application is a ϕ-value analysis, free from the adverse effects of mutations. These results will trigger the development of experimental techniques that enable the determination of accurate residue-specific equilibrium and kinetic parameters for analyzing the transition states of structural changes in proteins.

## Introduction

Protein molecules fold into their thermodynamically stable and biologically active structures[1]. This special property is acquired by evolution and optimized for each amino acid sequence. Particularly, small globular proteins fold quickly, typically in less than milliseconds, from unstructured states to their compact native structures. The entire process of protein folding is frequently a two-state transition without passing through any metastable intermediate states[2,3]. By increasing temporal and spatial resolutions, folding intermediates can be detected[4,5]. Nevertheless, the coarse-graining approximation of the folding process at the level of amino acid residues is useful for understanding the physicochemical basis of quick and smooth protein folding.

Protein folding can be described as an ensemble of the downslope motions of partially folded structures on a funnel-like energy landscape (Fig. 1A)[6–8]. The vertical axis represents the stability (free energy *G\**, which excludes the contributions from conformational entropy) of a partially folded structure, and the horizontal plane represents the conformational diversity (entropy *S\** originating from conformational entropy) of the ensemble. In the funnel view, a protein folds *via* multiple pathways. By contrast, the bimodal mass distributions obtained from hydrogen-deuterium exchange (HX) studies suggested that a protein folds along a defined pathway[9]. In the defined-pathway model, a protein molecule consists of smaller units, called ‘foldons’[10,11]. A foldon is a cooperative native-like structural element, and a folded foldon serves as a template for the folding of the next foldon, hierarchically. The folding pathway is predetermined by the same cooperative interactions that determine the final native structure. A recent reanalysis using the maximum entropy principle suggested that the multiple pathway model could explain the results of the HX study of cytochrome c[12]. However, the original experiment clearly showed the formation of one foldon (called blue) ahead of the other foldons, indicating the predefined formation of a foldon, at least in an early step of cytochrome c folding[9]. The funnel view *versus* the foldon view has been debated for decades[13,14]. Is it possible to resolve the conflict between the microscopic view (the smooth surface of a folding funnel) and the macroscopic view (the sequential formation of foldons)? Modeling of protein folding at a mid-level view is expected to be effective for this purpose[15]; for instance, at the residue level adopted in this study.

**Figure 1.**
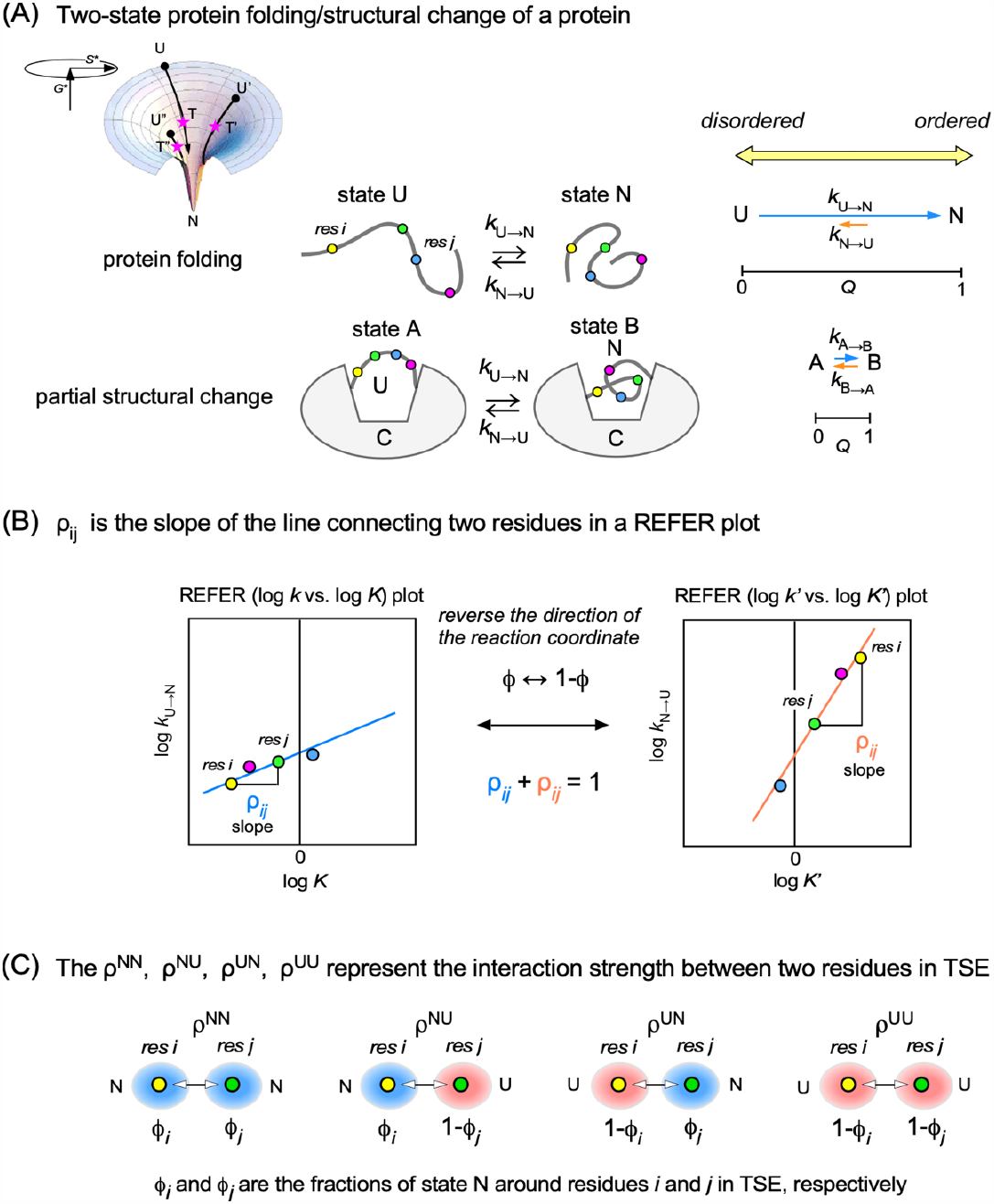
Basic aspects of the two-state structural exchange of a foldable polypeptide under the consistency principle of protein folding. (A) *Left*: funnel-like energy landscape. *Center*: the consistency principle was originally proposed for protein folding, but its application is now expanded to the smooth structural changes of proteins: a variable part exchanging between states U and N is embedded in a stationary constant region, C. *Right*: the structuredness of the states and the magnitude of the change from state U to state N vary depending on the systems. (B) The definition of the slope ρ_*ij*_ between residue *i* and residue *j* in a rate-equilibrium free energy relationship (REFER) plot, and the relationship between two slopes in two REFER plots, log *k* vs. log *K* and log *k*’ vs. log *K*’, where *k*=*k*_U→N_, *k’*=*k*_N→U_, *K*=*k*/*k*’, and *K*’=*k*’/*k*. (C) The ρ^NN^, ρ^NU^, ρ^UN^, and ρ^UU^ are constant coefficients introduced to calculate the slope ρ_*ij*_ between residue *i* and residue *j*. The state of a residue is specified by the fraction of state N around the residue in the TSE and expressed by the symbol ϕ.

What is the physicochemical basis for the conditions needed for a perfectly smooth energy landscape? Nobuhiro Gō proposed the ‘consistency principle’ theory in 1983[16]. He used a lattice model for simulation, in which each residue of a ‘protein’ is a point on a two-dimensional lattice and all pairs of residues in any partially folded structures have negative (i.e., preferable) short-range interaction energy *only if* the two residues are in contact within the native structure[17]. In other words, the consistency principle seeks the absence of non-native structures or interactions at any moment on the folding path from any random initial structure. The mathematical expression of the consistency principle can be defined as a special type of non-bonded contact interaction term, *V*=Σε_*ij*_, in a lattice Monte Carlo simulation[17] and a coarse-grained molecular dynamics simulation[18]:

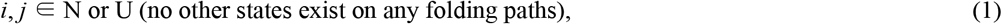

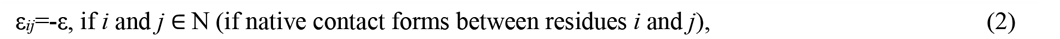

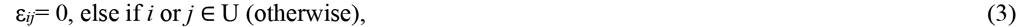

where ∈ means “being in the state of”, and N and U denote native and unfolded states, respectively. The solvent is not considered in these simulations. Gō and his colleagues showed that their simple lattice model can explain various physical properties of real proteins[16]. The 2D lattice model was subsequently extended to the cubic lattice model[19,20] and then to the coarse-grained Cα model[21]. These 3D version models had excellent predictive powers of folding transitions and pathways[22,23]. Multiple Monte Carlo 3D lattice model calculations revealed that a local maximum of free energy appears during protein folding. The significant decrease in the entropy term due to rapid structure formation resulting from the native-centric interactions is larger than the decrease in the enthalpy term, resulting in a barrier on the free-energy surface that corresponds to the transition state[24]. The consistency principle has been widely supported by molecular simulations, including the trajectory analysis of small proteins obtained by all-atom MD simulations with explicit water solvent[25]. On the assumption of the Gō model, the funnel-like energy landscape has a perfectly smooth surface (i.e., the range of microscopic roughness, *δE* = 0) so that it materializes the rapid folding of a polypeptide chain from any unfolded state. Although the absolute bias to the final folded state seems odd, it makes sense that the state realized by the consistency principle is equivalent to the ideal state of an actual gas. The ideal gas approximation is extremely useful as a reference for understanding real gases.

Bryngelson and Wolynes proposed a more realistic version of the protein folding principle in 1987 (i.e., minimal frustration model), based on the theory of spin glasses[26]. Real proteins have rugged energy surfaces because frustration occurs owing to the impossibility of satisfying all favorable energetic interactions simultaneously, and results in a rough energy landscape surface[27]. To be kinetically foldable, the funnel must have a sufficiently steep slope so it can overcome local traps created by the frustration. Through evolutionary selections, the roughness of the rugged surface of the funnel-like landscape is minimized (*δE* < 3*k*_*B*_*T*) by balancing between rapid folding rates and functionality requirements, such as in catalytic and allosteric proteins[28,29].

NMR spectroscopy can probe the local environment of a polypeptide chain. This feature enables NMR to determine the residue-specific equilibrium constant, *K*, and the residue-specific rate constants, *k* (forward) and *k*’ (backward), of the two-state exchange of a polypeptide chain. Unexpectedly, we found that the observed constants varied from one residue to another in the two-state topological isomerization of a 27-residue bioactive peptide, nukacin ISK-1[30,31]. Such variations in the thermodynamic and kinetic parameters of a polypeptide chain have occasionally been reported over the years[32,33] and global fitting is often applied to obtain the average values, but the significance of the variations has remained enigmatic. This is probably because the artifacts caused by systematic measurement biases and random measurement errors cannot be ruled out. Intriguingly, we found a linear relationship between the residue-specific log *K* values and the residue-specific log *k* (also log *k*’) values for nukacin ISK-1, even after eliminating biases and errors from the measurement and analytical procedures for determining these parameters[34]. This linear relationship must be more than a coincidence.

The double logarithmic relation of the two parameters, *K* and *k*, is generally called REFER (rate-equilibrium free energy relationship) (Fig. 1B), and the linear relationship in the REFER plot is referred to as LFER (linear free energy relationship), since the two axes, log *K* and log *k*, are proportional to the changes in the corresponding free energy terms. In particular, the LFER we found is referred to as residue-based LFER (rbLFER) because it is derived from amino acid residues in one polypeptide chain[31]. To test the general applicability of rbLEFR, we performed a retrospective study by collecting residue-specific equilibrium and rate constants in exchange processes, including protein conformational changes, the coupled folding and binding of intrinsically disordered polypeptides (IDP), and the structural fluctuations of folded globular proteins. We reanalyzed the data in the original reports by creating 18 REFER plots. Among them, we found 10 that seem genuine [35]. Since no reports assumed a relationship between log *K* and log *k*, it is doubtful that intensive efforts were made to reduce the measurement biases and errors in the previous studies, except for our nukacin ISK-1 case[34]. Considering the errors left unintentionally, we concluded that the rbLFER holds in a wide variety of protein-related phenomena[35].

As mentioned above, the consistency principle is a physical model of ideal foldable polypeptide chains. We previously suggested that the rbLFER was an experimental consequence of the consistency principle[31]. Here, we examined the theoretical relationship between log *K* and log *k* under the consistency principle (Fig. 2). We found that the alignment of data points on a parabolic curve in a REFER plot is the general solution (Fig. 2B). The mathematical expression, referred to as the quadratic free energy relationship (QFER), enabled us to devise a method for analyzing the transition states in the structural changes of proteins (Fig. 2C and 2D).

**Figure 2.**
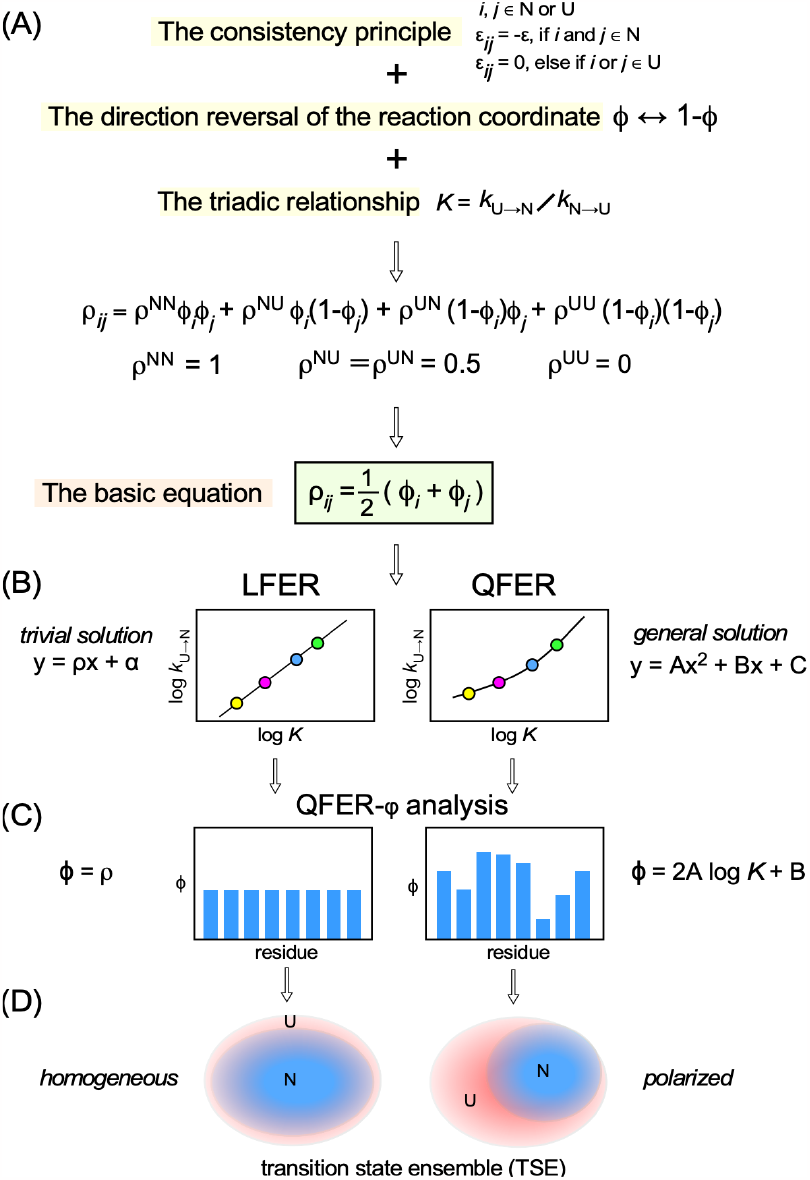
Outline of the transition-state analysis based on the consistency principle of protein folding. (A) The combination of the consistency principle, the direction reversal of the reaction coordinate, and the triadic relationship, *K*=*k*/*k*’, leads to the basic equation expressing the relationship between ρ_*ij*_ and ϕ_*i*_/ϕ_*j*_. (B) The linear and quadratic relationships in the REFER plot are derived from the basic equation. (C) The QFER-φ analysis provides per-residue ϕ values. (D) A homogeneous TSE is formulated by residue-based LFER (linear free energy relationship), and a polarized TSE is formulated by residue-based QFER (quadratic free energy relationship).

## Methods

Quadratic OLS (ordinary least squares) fitting was performed with the standard polynomial fitting in Excel, and quadratic TLS (total least squares) fitting was performed by solving cubic equations (Appendix I) with the add-in program, Solver, in Excel. The protein cartoons were generated with the program PyMOL, version 2.4.2 (Schrödinger).

The *F*-statistic was used to assess the improvement in fit by using QFER relative to LFER. The *F*-statistic is defined as:

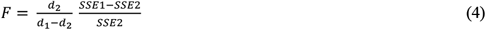

where *SSE*1 and *SSE*2 are the sum squared-error residuals for the LFER and QFER models, respectively, and *d*_1_ and *d*_2_ are the degrees of freedom (*d*_1_ > *d*_2_), respectively. The *F*-statistic calculated from the experimental data is compared with the (1-α)100% critical value of the *F*-statistic obtained from the theoretical distribution of *F*_(*d1*-*d2*),*d2*_, under the assumption of a normal distribution of the experimental data. α is the significance level, which defines the sensitivity of the *F* test.

The jackknife (i.e., leave-one-out) method was used to estimate the ϕ-values. The partial estimates are a set of the slopes (=2Alog K + B) of the quadratic curves in REFER plots obtained by iteratively performing quadratic OLS fits after leaving out one data point from the data set. The TLS fitting is better, but we used the OLS fitting as a convenient alternative. The jackknife ϕ-value was determined as the average of the partial estimates of ϕ. The standard error of the jackknife ϕ-value was computed by multiplying the standard error of the partial estimates of ϕ by *n*-1, where *n* is the number of data points[36].

## Results

The consistency principle is widely applicable to various protein-related processes, in addition to protein folding[37]. For general structural changes, the consistency principle can be applied to a specific part surrounded by a constant region (Fig. 1A). The specific part becomes a structured state from a less structured state (which may not be a random structure). The structural difference between the initial and final states of the specific part is equivalent to the native structure formation in the Gō model. Although the experimental data used as an illustrative example in the present study are not those of protein folding, we will use states U and N, instead of states A and B, to maintain the essence of the native-centric concept.

### The slope in a REFER plot is related to the free energy changes of reaction and activation

Let ρ_*ij*_ denote the slope of the line segment connecting two points corresponding to residues *i* and *j* in a REFER plot (Fig. 1B):

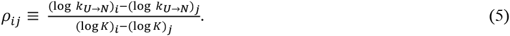

An equilibrium constant is related to the standard Gibbs free energy change of reaction Δ*G°* by

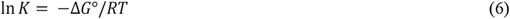

and a rate constant is related to the standard Gibbs free energy change of activation Δ*G*^‡^ by

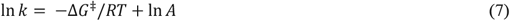

where *R* is the universal gas constant, *T* is the absolute temperature, and *A* is the pre-exponential factor. In the Arrhenius equation, *A* is a temperature-independent constant, and in the Eyring equation, *A* = *κk*_*B*_*T/h*, where *κ* is the transmission coefficient, *k*_B_ is the Boltzmann constant, and *h* is the Planck constant. Eq. 5 is rearranged by using Eqs. 6 and 7:

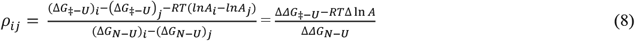

where the symbol ΔΔ denotes the difference between the two energy terms, and the subscripts *i* and *j* are omitted for simplicity. See Fig. 3 for the definitions of ΔG_‡-U_ and ΔG_N-U_. We assume a constant pre-exponential factor value among different residues; i.e., Δln*A* = 0. The rational explanation is that the parameters that determine non-bonded interactions should be identical for all residues, which leads to the identical transmission coefficient *κ* in the Eyring equation for all residues within a polypeptide chain. The same assumption was adopted in previous protein folding studies, in which the exchange processes were perturbed by solvent condition changes and amino acid substitutions[38,39].

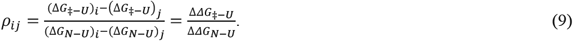

**Figure 3.**
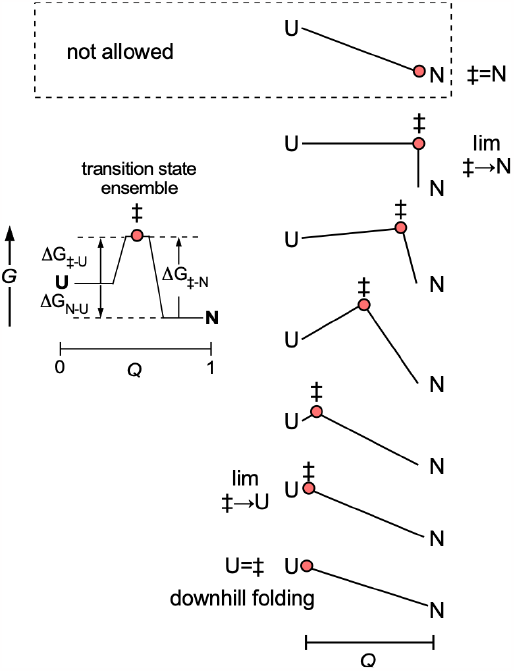
Free energy level diagram with varied positions of the transition state ‡ on the reaction coordinate *Q*. The red-filled circles indicate the position of the transition state ensemble on the reaction coordinate. The *lim* ‡→N and *lim* ‡→U denote the limit as the transition state ‡ approaches state N and state U, respectively.

### Calculation of the slope in the REFER plot under the consistency principle

We start by considering the transition state ensemble (TSE) generated by starting from all initial states. We assume that the slope ρ_*ij*_ between two data points corresponding to two residues *i* and *j* in a REFER (log *k*_U→N_ vs. log *K*) plot can be calculated as an average of four coefficients, ρ^NN^, ρ^NU^, ρ^UN^, and ρ^UU^, by weighting with the fractions of the local states around residues *i* and *j* in the TSE. These coefficients are constants representing the “interaction strength” in the TSE between two residues (Fig. 1C). The strict definitions of the coefficients will be provided below. Next, the state of residue *i* is specified by the fraction of state N around the residue in the TSE and expressed by the symbol ϕ_*i*_. The consistency principle assumes that the TSE only contains residues in either state N or state U (Eq. 1). Consequently, 1-ϕ_*i*_ denotes the fraction of state U of residue *i* in the TSE. The product ϕ_*i*_ϕ_*j*_ represents the fraction of the coexisting N states around residues *i* and *j*, and so on.

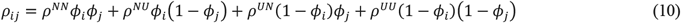

We first evaluate the value of ρ^UU^. When setting ϕ_*i*_=ϕ_*j*_=0 in Eq. 10, ρ_*ij*_ = ρ^UU^. In the two-state exchange folding process, the transition state can approach and match state U exactly (Fig. 3). In the latter extreme condition, the overall folding process is a so-called downhill process [40]. The direct substitution of U for ‡ in Eq. 9 results in

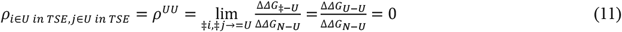

where ‡→= U means that the transition state approaches and finally matches state U.

### Definitions of ρ^NN^, ρ^NU^, and ρ^UN^

When setting ϕ_*i*_ = ϕ_*j*_ =1 in Eq. 10, then ρ_*ij*_ = ρ^NN^. The direct substitution of N for ‡ in Eq. 9 results in

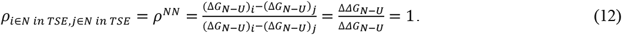

Similarly, setting ϕ_*i*_ = 1 and ϕ_*j*_ = 0 in Eq. 10 leads to ρ_*ij*_ = ρ^NU^, and setting ϕ_*i*_ = 0 and ϕ_*j*_ = 1 leads to ρ_*ij*_ = ρ^UN^. Direct substitutions of N and U for ‡ in Eq. 9 result in

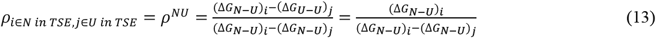

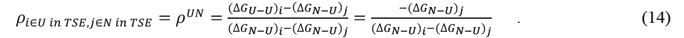

The addition of Eq. 13 and Eq. 14 provides the expression, ρ^NU^ + ρ^UN^ = 1. These results are correct, and the operations seem valid, but they are not allowed because they violate the definition of the two-state exchange. In the two-state exchange, the absence of any intermediate states applies the highest free energy level of the transition state throughout the reaction coordinate (Fig. 3). Consequently, the transition state can move from state U to a position very close to state N, but the position of the transition state cannot match state N exactly (the ‡=N case in Fig. 3).

To be exact, we must consider the limiting case of ‡→N for the proper definitions of ρ^NN^, ρ^NU^, and ρ^NU^.

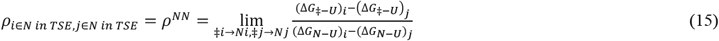

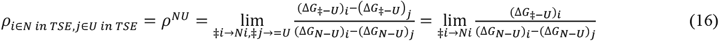

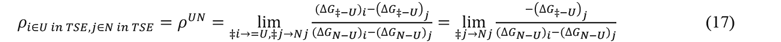

It is tempting to separately evaluate the individual terms that constitute the definition formulae (Eqs. 15-17) and then perform arithmetic operations to find the values of *ρ*^NN^, *ρ*^NU^, and *ρ*^UN^. Under the assumption of the consistency principle, however, the free energy values of the transition state G_‡_ and those of the two ground states, G_U_ and G_N_, are strongly correlated, and consequently the (ΔG_‡-U_)_*i*_, (ΔG_‡-U_)_*j*_, (ΔG_N-U_)_*i*_, and (ΔG_N-U_)_*j*_ terms are intercorrelated to each other. Thus, we will deduce the values of *ρ*^NN^, *ρ*^NU^, and *ρ*^UN^ from the consistency principle in combination with obvious premises in the next section.

### The combination of the reverse direction of the reaction coordinate with the triadic relationship of *K, k*, and *k*’ constrains the relationships between ρ^NN^, ρ^NU^, ρ^UN^, and ρ^UU^

The slope in the REFER (log *k*_U→N_ vs. log *K*) plot is calculated using Eq. 10. For convenience, Eq. 10 is redisplayed with some details.

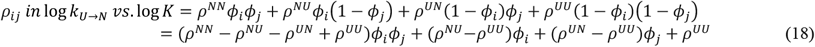

Now we will view the same folding process from the reverse direction. The slope in another REFER (log *k*_N→U_ vs. log *K*’) plot can be calculated by replacing ϕ with 1-ϕ in Eq. 18 (Fig. 1B).

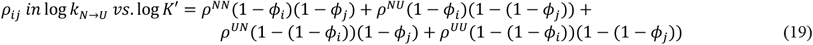

where *K*’ = 1/*K* or log *K’* = -log *K*. After rearrangement,

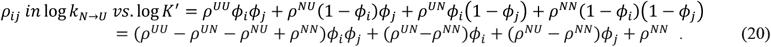

From the relationship *K* = *k*_U→N_ /*k*_N→U_, we obtain log *k*_U→N_ -log *k*_N→U_ = log *K*. Differentiating both sides with respect to log *K* gives

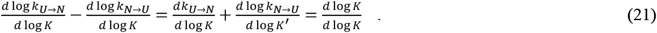

The right two terms of Eq. 21 are equivalent to

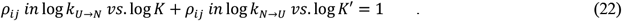

Substituting Eqs. 18 and 20 into Eq. 22, we obtain:

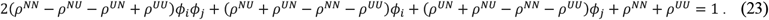

This equation must hold for any values of ϕ_*i*_ and ϕ_*j*_, which results in the following system of equations:

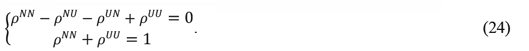

The substitution of ρ^UU^ = 0 (Eq. 11) into the simultaneous equations provides

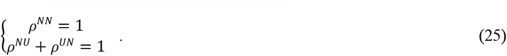

ρ^NU^ and ρ^UN^ are the slopes connecting two residues, each in different states, in the TSE in a REFER plot. In the consistency principle, non-bonded interactions that facilitate the structure formation are position-independent (Eq. 2), and therefore it is reasonable to assume ρ^NU^ = ρ^UN^. Another explanation for the basis of this assumption will be provided later.

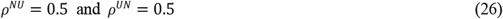

In summary, the combination of the consistency principle (Eqs. 1-3), the reverse view of the folding process (Eq. 20), and the triadic relationship between the equilibrium and the two rate constants (Eq. 21) derives ρ^NN^ = 1 and ρ^NU^ = ρ^UN^ = 0.5 from ρ^UU^ = 0 (Fig. 2). The assumptions are a constant pre-exponential factor *A* among residues in a polypeptide chain (Eq. 8) and the mathematical formula of the slope ρ_*ij*_ using the four coefficients, ρ^NN^, ρ^NU^, ρ^UN^ and ρ^UU^ and the ϕ values around residues *i* and *j* in TSE (Eq. 10).

### Physicochemical insight from ρ^NN^ = 1

First, we transform Eq. 9 as follows:

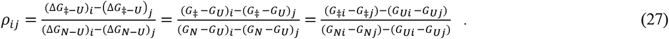

As mentioned, we define ρ^NN^, ρ^NU^, and ρ^NU^ as the limits of the slope ρ_*ij*_ as the transition state approaches state N.

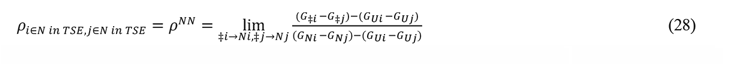

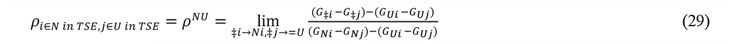

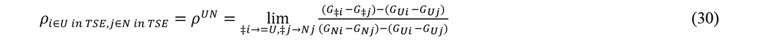

The ‡→= U condition results in

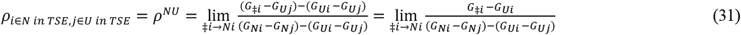

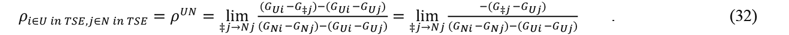

The addition of Eqs. 31 and 32 yields

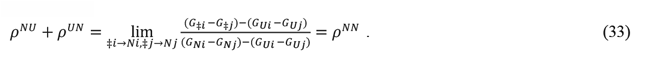

Since the value of ρ^NN^ is 1, Eq. 28 provides

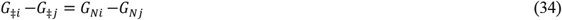

in the limit of ‡→N around residues *i* and *j*, and Eq. 33 reproduces Eq. 25.

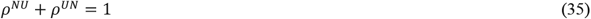

Eq. 34 means that the free energy difference between two residues in the TSE (G_‡*i*_ - G_‡*j*_) is equal to the free energy difference between the same residues in state N (G_N*i*_ - G_Nj_), demonstrating the structural and interaction similarities between the transition and native states in the limit of ‡→N under the consistency principle. Note that the free energy level of individual residues in the TSE is always higher than that in the native state (the ‡→N case in Fig. 3), reflecting the structure and interactions specific to the TSE. The structural identity shared between the transition state and the final state can be extracted and is usable only after viewing the TSE from the difference in the residue-specific free energy levels of two residues.

### Arithmetic relation between ρ and ϕ

The assignment of the ρ^NN^, ρ^NU^, ρ^UN^ and ρ^UU^ values within Eq. 10 gives:

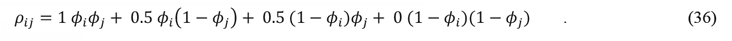

This formula is simplified to

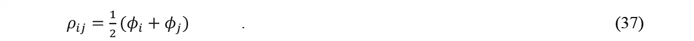

This equation means that the slope of the line segment connecting residues *i* and *j* in a REFER plot is equal to the average of the ϕ values of the same pair of residues. Let *n* be the number of observed residues. The number of ϕ is *n*, and the number of ρ is ½*n*(*n*-1). If Eq. 37 is valid, then the average and variance of ρ_*ij*_ are calculated to be

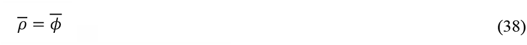

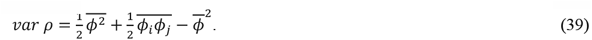

Refer to Appendix II for the derivation.

If all data points lie perfectly on the linear line in a REFER plot, then all ρ_*ij*_ have the same value. Let this value be ρ.

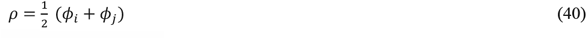

The solution of Eq. 40 is the single value ϕ for all residues,

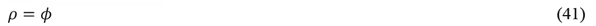

which means that all residues share a single ϕ value in the TSE. In other words, the TSE is homogenous in the case of rbLFER.

### Slope ρ indicates the TSE position on the reaction coordinate

The reaction coordinate of protein folding is generally defined as the global structural measures of similarity to the native state, such as the fraction of native contacts (*Q*), the similarity of natively contacting residue pairs to their native distances (*Q*_S_), the average shortest path length (<*L*>), and the radius of gyration (*R*_g_)[41]. Given the free energy profile of the folding landscape using any of the structural reaction coordinates, ϕ determines the position of the transition state on the reaction coordinate[41]. Based on the relationships, ρ = ϕ (Eq. 41) in the case of homogeneous TSE and 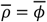 (Eq. 38) in the case of polarized TSE, the slope ρ indicates the position of the transition state on the native structure-based reaction coordinates. In other words, the slope ρ represents the structural similarity between the transition state and the final state. Consequently, the slope ρ has a value between 0 and 1, inclusive.

Given the physical meaning of the slope ρ, another interpretation of the values of ρ^NN^, ρ^NU^, and ρ^NU^ is possible. When two residues in the TSE are both in state N (ρ = ρ^NN^), the two residues behave as if the folding process proceeded to the end on the reaction coordinate (*φ*= 1). Thus, *φ*= ρ^NN^ = 1. When one residue is in state N and the other residue is in state U in the TSE (ρ = ρ^NU^ or ρ^UN^), it is reasonable to assume that the folding process proceeds halfway along the reaction coordinate (*φ*= 0.5). Therefore, the assignment of *φ*= ρ^NU^ = ρ^UN^ = 0.5 is understandable.

### QFER is the general solution

Let the equation of the solution curve be log *k* = *f*(log *K*) in a REFER plot under the restriction of the basic equation, Eq. 38. The use of the function *f* assumes that the kinetical properties of a two-state exchange (i.e., log *k*_U→N_, log *k*_N→U_, ϕ_*i*_, and ϕ_*j*_) are uniquely determined if the equilibrium property (i.e., log *K*) is fixed. We use x as a replacement for log *K* for simplicity. The next equation is obtained from Eq. 37.

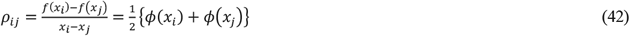

Here, we aim to find an *ideal* solution curve on which *all pairs* of points satisfy the basic equation. This requirement condition allows the limiting operation of x_j_ → x_i_ to find an ideal solution curve:

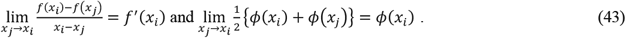

Thus, ϕ is the derivative of *f*(x).

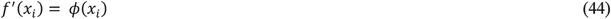

By substitution, we obtain the following.

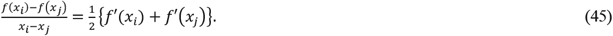

Assume that x_*j*_ = 0 and x = x_*i*_ without loss of generality.

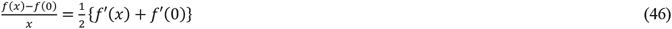

Set C = *f*(0) and B = *f’*(0), and y = *f*(x) – C.

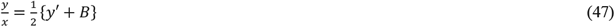

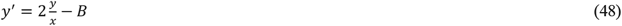

By setting *u* = y/x, we have

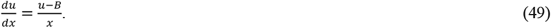

By integrating Eq. 49, yet another constant A is introduced. The solution is a parabolic curve, *f*(x) = Ax^2^ + Bx + C. Refer to Appendix III for the derivation. The assignments of log *k* and log *K* to this formula provide the following quadratic free energy relationship (QFER).

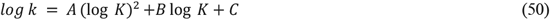

Here, ϕ is equal to the slope of the tangent at the data point (Eq. 44).

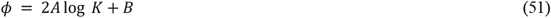

Eq. 51 indicates that the consistency principle is compatible with a polarized TSE, in which ϕ varies from residue to residue.

We now describe the relationships among the coefficients of the regression equations. In the case of LFER, we obtain a linear formula, log *k* = ρ log *K* + α. Then, the assignment of the triadic relation, log *k’* = log *k* – log *K*, results in another linear formula, log *k’* = (ρ-1) log *K* + α. Thus, the difference between the slopes of the two linear formulas is always 1, and the coefficients of the constant terms are the same. In the case of QFER, we obtain a quadratic formula, log *k* =A (log *K*)^2^ + B log *K* + C. The assignment of log *k’* = log *k* – log *K* then results in log *k’* = A(log *K*)^2^ + (B-1) log *K* + C. Thus, the difference between the coefficients of the linear term is always 1, and the coefficients of the squared term and the constant term are the same. Lastly, the sign of the squared term, A, must be positive to make ϕ physically meaningful, because the fraction of state N in the TSE (i.e., ϕ) should increase simultaneously with the fraction of state N in the ground state ensemble (GSE) (i.e., increased state N fraction in GSE means increased log *K*), as clarified by Eq. 51.

The following discussion should provide useful clarifications for readers. Under the assumption of the consistency principle, no non-bonded interactions work between two amino acid residues if one of the residues is in state U (Eq. 3). It is tempting to assume ρ^NU^ = ρ^UN^ = 0, in addition to ρ^NN^ = 1 and ρ^UU^ = 0, which results in the likely basic equation,

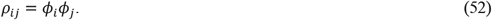

It is possible to solve a differential equation,

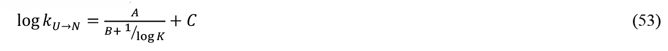

where A, B, are C are arbitrary constants. According to the triadic relationship of log *K* = log *k* – log *k’*, we derive

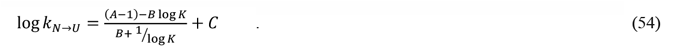

The resultant functional form of log *k*_N→U_ in terms of log *K* is not the same as that of log *k*_N→U_. This violates the necessary condition that the two directional views of the same exchange phenomenon must be described by the same functional form in terms of log *K*.

In summary, although ρ^NU^ = ρ^UN^ = ρ^UU^ = 0 seems appealing, it cannot provide the desired general solution. Eq. 3 describes the contribution of the interaction of a pair of residues to the enthalpy term of the system. To evaluate the ρ^NU^, ρ^UN^, and ρ^UU^ values properly, we must consider the entropy term. When one residue is in state N in the TSE, the structure formation around the residue affects the entropy of the system. This is the reason for ρ^NU^ ≠ ρ^UU^ and ρ^UN^ ≠ ρ^UU^.

### Comparison between the OLS and TLS fittings

Curve fitting is generally performed with the ordinary least squares (OLS) method. OLS assumes errors in the measurement of log *k* (i.e., y-axis) and no errors in that of log *K* (x-axis). OLS minimizes the sum of the squares of the vertical offsets to determine the fitting curve shape (Fig. 4). However, errors are also present in the measurement of log *K*. The total least squares (TLS) method is appropriate in this case. TLS minimizes the sum of the squares of the perpendicular offsets, instead of the vertical offsets. Refer to Appendix I for the details of the procedure. Practically, however, the resultant quadratic curves are not substantially different (Fig. 4). Thus, we use the results of the OLS fitting hereafter.

**Figure 4.**
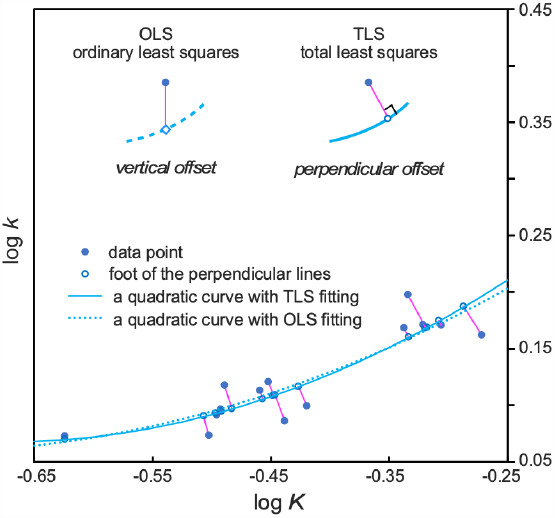
Comparison of two regression methods, OLS (ordinary least squares) and TLS (total least squares). The *insets* show the differences in the definition of an offset (i.e., the lengths of the magenta line segments). OLS and TLS seek the regression curve parameters that minimize the sum of the squares of the offsets.

### Nukacin ISK-1 system as an illustration

In our previous retrospective analysis, we interpreted the REFER plots on the assumption of LFER[35]. Here, we reinterpreted the same REFER plots using QFER. In most instances, the fitting did not improve, and we must reluctantly accept that the TSE is homogeneous, considering the large, non-negligible measurement errors. As exceptions, curve fitting improved in two cases: the two-state topological interconversion of nukacin ISK-1 and the coupled folding and binding of the pKID peptide to the KIX domain. Below, we will use the reanalysis of nukacin ISK-1 as an example. Nukacin ISK-1 is a 27-residue lantibiotic peptide (Fig. 5)[42]. The two sets of ^1^H–^15^N amide cross peaks in the ^1^H–^15^N HSQC (heteronuclear single quantum coherence) spectrum indicated the existence of two states in solution, termed states A and B[43]. The interconversion rate was slow on the sub-second time scale, reflecting the high energy barrier, 10*k*_*B*_T[30,44] (See Appendix IV for the history of corrections). We found residue-based linear relationships between log *k*_A→B_ and log *K* and between log *k*_B→A_ and log *K* in the REFER plot[31,34]. Because the structures of the two states have different backbone topologies, some side chains are threaded through the main chain ring formed by one of the three mono-sulfide linkages.

**Figure 5.**
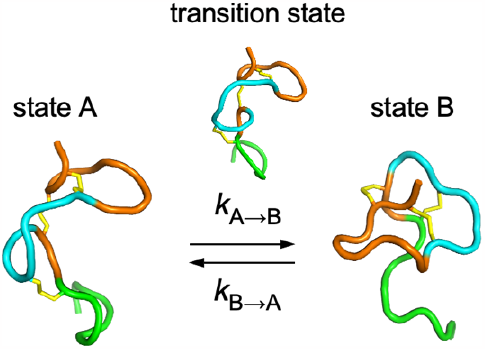
The two-state topological interconversion between states A and B of nukacin ISK-1. The amino acid sequence is color-coded according to the N-tail (green), hinge (cyan), and core (orange) regions. Nukacin ISK-1 contains three mono-sulfide linkages (yellow). The 3D structures of states A and B were determined by NMR [PDB IDs: <u>5Z5Q</u> and <u>5Z5R</u>, respectively]. The 3D structure of the transition state was inferred from the targeted MD simulation[30].

### Comparison between LFER and QFER

Figure 6 shows the fittings of nukacin ISK-1 data with LFER and QFER. The R^2^ values are higher in the QFER fitting than the LFER fitting. We performed the *F*-test to assess whether the improvement in fit obtained by applying QFER is significant, or merely arises from the random statistical reduction in the *SSE* (sum squared-error residual) due to an additional parameter. The *F*-statistic was calculated according to Eq. 4 and compared with the critical value obtained from the theoretical *F* distribution. Unfortunately, the *F*-test revealed that the improved fittings with QFER (i.e., larger R^2^ values) were *not* statistically significant (Appendix V). In our nukacin ISK-1 case, we think that there is still room to improve the peak volume determination of the EXSY NMR spectra.

**Figure 6.**
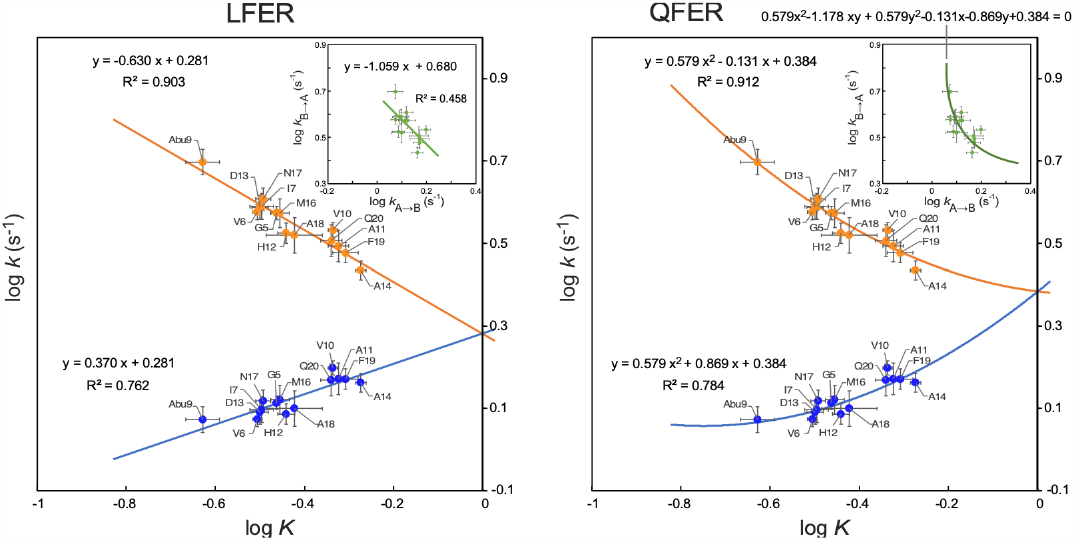
Comparison of LFER and QFER analyses of nukacin ISK-1. The blue line/curve represents the log *k*_A→B_ vs. log *K* plot, and the orange line/curve represents the log *k*_B→A_ vs. log *K* plot. The *insets* show the log *k* vs. log *k’* (log *k*_B→A_ vs. log *k*_A→B_) plots with a fitted line or a fitted quadratic curve.

As discussed previously[35], the triadic relationship, *K* = *k*/*k’* or log *K* = log *k* – log *k’*, could generate an artificial linear relationship in the REFER plot if a careless experimenter generated a REFER plot with unignorable measurement biases and/or large measurement errors. To identify a true rbLFER/rbQFER, the correlation between log *k* and log *k’* must be evaluated (*insets*, Fig. 6)[35]. The log *k* vs. log *k’* plot must have a negative correlation to make rbLFER/rbQFER physically meaningful. In the case of LFER, log *k* vs. log *k’* has a linear negative relationship. In the case of QFER, log *k* vs. log *k’* has a tilted parabolic relationship (Appendix VI).

### Polarity and polar pattern of the TSE

We debated the local structural identity between the transition state and the native states under the consistency principle. To describe the structural identity quantitatively, we formally define an apparent equilibrium constant *K*^‡^ on a per-residue basis, as follows:

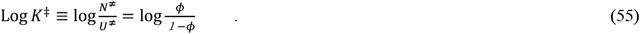

To be precise, an equilibrium constant cannot be defined in a transient process. In some special situations, however, the treatment of a quasi-equilibrium is useful in transient state analyses. For example, in the transition state theory, *K*^‡^ is defined as [AB]^‡^/[A][B] in the reaction of A+B ⇄ AB^‡^ → P, which successfully led to the Eyring equation. By substituting Eq. 51 into Eq. 55, we obtain the following:

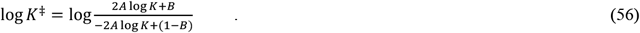

Figure 7 shows the curve defined by Eq. 56 (red dotted curve) and the approximate straight-line segment (black line) in the ϕ range of 0.1 to 0.9. The line segment is expressed as follows:

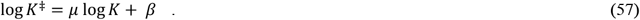

**Figure 7.**
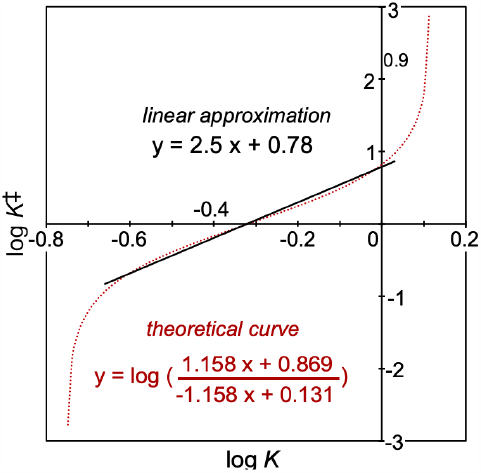
Log *K*^‡^ vs. log *K* plot of nukacin ISK-1 The red dotted line is the theoretical curve defined by Eq. 56. The black solid line is the linear approximation (Eq. 57) in the ϕ range from 0.1 to 0.9.

This is LFER between log *K*^‡^ and log *K*. The slope can be approximated as μ ≈ 4A (Appendix VII).

We define ‘polarity’ as the distribution of the *K*^‡^ values in the TSE state, and ‘polar pattern’ as the spatial distribution of the *K*^‡^ values in the TSE structure. Similarly, we define the polarity and polar pattern of GSE using the *K* values. Eq. 57 indicates that the polar patterns of TSE and GSE are closely interrelated, under the assumption of the consistency principle. The slope μ is an indicator for the polarization degree of TSE relative to GSE. If μ is zero, then TSE has no connection to GSE. The REFER plot is described by LFER (A = 0), and the TSE is homogenous. Next, if 0 < μ < 1, then TSE is slightly polar. If μ ≥ 1, then TSE is more polar than GSE. In the case of nukacin ISK-1, μ is equal to 2.5 (Fig. 7). In the transition state, some side chains are threaded through the main chain ring formed by one of the three mono-sulfide linkages (Fig. 5). This may be the reason for the higher polarization degree of TSE than that of GSE.

### Structural insights into the transition states

The consistency principle now expands its application to the smooth structural changes of proteins[37], with various structuredness of the states and magnitudes of the changes (Fig. 1A). Nukacin ISK-1 has a small hydrophobic core around residue Phe21 in state B[30], and hence the state B structure is more compact than the state A structure (Fig. 5). The change in the direction of A→B can be regarded as a protein folding process, U→N, of a small portion around Phe21. The variable region around Phe21 is embedded in an invariant structured region. Thus, the REFER plots in Fig. 6 provide information about the transition state in the direction of structure formation (blue lines, *k*_A→B_).

A visual inspection reveals some clusters of the (log *K*, log *k*) data points. For clarification, the data points are color-coded according to the structures of the destination states; i.e., state B for nukacin ISK-1. As shown in Fig. 5, the amino acid sequence of nukacin ISK-1 is divided into three regions, N-tail, hinge, and core, by reference to the state B structure[30]. Residues with the same color tend to form clusters in the REFER plot (Fig. 8). This suggests that the adjacent residues in the same cluster change the states in a coordinated manner. Thus, the clusters identified in the REFER plot agree with the foldon concept.

**Figure 8.**
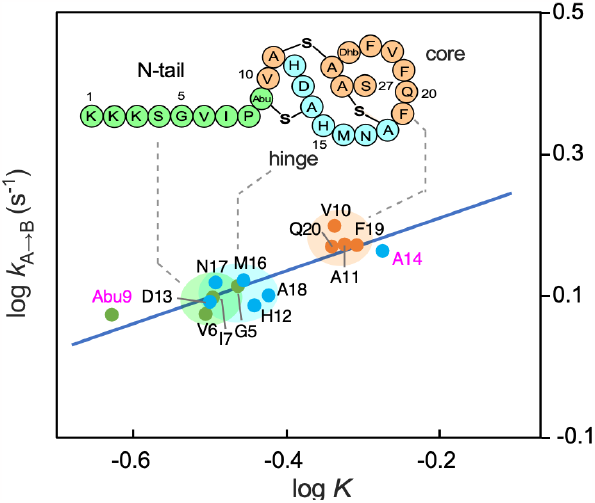
Data points form a couple of clusters in the REFER plot of nukacin ISK-1. The cluster formation reveals the connection between the energetic view and the structural view. The energetic view is the cluster formation of the (log *K*, log *k*) data points, whereas the structural view is the color-coded segmentation of the amino acid sequences by reference to the three-dimensional structure. The magenta-highlighted text marks the exceptional residues, which are on the lines/curves but away from the clusters to which they belong.

There are exceptional residues, which are defined as the residues that are far from the clusters to which they belong: Abu9 and Ala14 in nukacin ISK-1 (magenta, Fig. 8). The side chains of Abu9 and Ala14 of nukacin ISK-1 are linked by a mono-sulfide bond and involved in the 6-membered main-chain ring. As discussed in detail previously[34], Abu9 and Ala14 probably became exceptions by adopting special states in the TSE to enable the threading of the His12 and Asp13 side chains through another main-chain ring.

### QFER-φ analysis

The ϕ-value analysis based on Eq. 51 is referred to as a QFER-φ analysis. We use a variant character φ, in place of ϕ, to consider the possibility of different ϕ values obtained by various methods. When we apply the QFER-φ analysis to the structural changes of a small portion surrounded by a constant region (Fig. 1A), φ is the ϕ value of the small portion. The ϕ-value for a whole protein molecule is calculated by

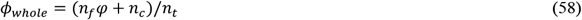

where *n*_f_ is the residue number involved in the focused small portion, *n*_c_ is the residue number of the constant region, and *n*_t_ (= *n*_f_ + *n*_c_) is the total residue number. The φ and ϕ_whole_ values become equal in protein folding studies (*n*_c_ = 0). Thus, the φ-value defined in this study is different from the popular ϕ-value obtainable from the mutational Φ analysis specialized for protein folding studies [38,45–47].

The QFER-φ analysis provides information about the polarized structure formation in the TSE state. Eq. 51 indicates that the ϕ value is the slope of the tangent at the data point corresponding to the residue. Because a data point is not always on the curve due to measurement errors, the nearest point on the curve found by the TLS fitting or conveniently by OLS fitting (Fig. 4) is used to calculate the slope of the tangent. We used the jackknife method to estimate the φ values and their standard errors (Fig. 9).

**Figure 9.**
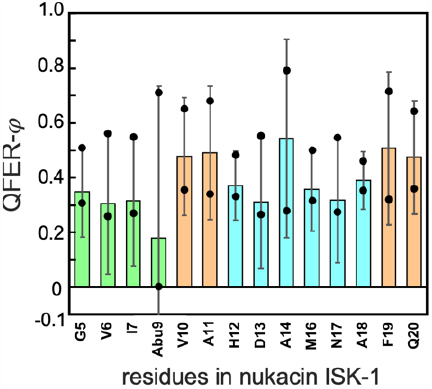
QFER-φ analysis. The partial estimates of the ϕ-values in the two-state exchanges of nukacin ISK-1 were calculated by the jackknife method. The averages of the partial ϕ-value estimates are displayed as bar graphs with the standard errors. The minimum and maximum values of the jackknife partial estimates of the ϕ-values are designated by black dots. The color coding is the same as in Fig. 8.

## Discussion

In our previous NMR study, we found that the two-state topological isomerization of a 27-residue peptide, nukacin ISK-1, had large distributions of the residue-specific equilibrium constant, *K*, and the residue-specific rate constant, *k*, which implied the reduced degrees of thermodynamic and kinetic cooperativities[30]. Interestingly, the log *k* vs. log *K* plots showed a linear free energy relationship (LFER)[31]. This discovery prompted us to collect residue-specific equilibrium and residue-specific rate constants from published sources, to analyze a substantial number of residue-based LFERs in a wide variety of protein structural changes[35]. In this study, we derived the basic equation 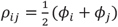 (Eq. 37), from the consistency principle of protein folding (Eqs. 1-3). In the derivation, we adopted three simple and reasonable assumptions: a constant pre-exponential factor *A* among residues in a polypeptide chain (Eq. 8), the mathematical formula of the slope ρ_*ij*_ using the four ρ coefficients and the ϕ values around residues *i* and *j* in the TSE (Eq. 10), and log *k* and ϕ are uniquely determined as a function of log *K* (Eq. 42). The equality of ρ^NU^ and ρ^UN^ (Eq. 26) might be arguable, but it is more unnatural to assume that they have different values, because non-bonded interactions that facilitate the structure formation are position-independent under the consistency principle (Eq. 2).

The consistency principle was originally proposed for the smooth folding of foldable proteins (i.e., naturally selected amino acid sequences) from completely unstructured states[16], and is now applicable to protein-related phenomena including conformational changes, native-state dynamics, and ligand binding[37]. For the cases of conformational changes, the consistency principle applies to a small part surrounded by a constant region (Fig. 1A). The conformational changes of the small part from a certain state (which may not be a random structure) to a more structured state are regarded as the protein folding process that follows the Gō model. The basic equation 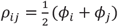 specifies the relationship between the slope ρ in the REFER (i.e., log *k* vs. log *K*) plot and the residue-specific ϕ values (i.e., the fraction of the destination state in the transition state ensemble). We derived an *ideal* solution curve on which all pairs of points satisfy the basic equation. The (log *K*, log *k*) data points lie on a straight line if the transition state ensemble (TSE) is homogeneous and on a parabolic curve if the TSE is polarized. We refer to the linear relationship as residue-based LFER and the parabolic relationship as residue-based QFER (Fig. 2). rbLFER/rbQFER provides a framework for analyzing the transition state in two-state structural exchanges of proteins. The first application is the elucidation of a linear relationship between log *K*^‡^ and log *K* (Fig. 7). The slope μ is an indicator of the polarization degree of TSE relative to GSE and can be approximated as μ ≈ 4A. If A = 0, then QFER reduces to LFER, and the TSE state is homogenous. The second application is the QFER-φ analysis (Figs. 2C and 9). The ϕ value is obtainable as the slope of the tangent nearest the data point, using Eq. 51.

Among the 18 REFER plots we made in the previous study[35], 10 showed residue-based LFER. Without considering the possible relationship between log *K* and log *k*, it is easy to imagine that the efforts to reduce measurement biases and errors were inadequate. In the presence of non-negligible biases and errors, even if the rbQFER is true, the relationship between the log *K* and log *k* values can only be analyzed with rbLFER. In this study, we chose the slow-state exchange of a bioactive peptide, nukacin ISK-1 (Fig. 5), as an example to test the feasibility of the residue-based QFER. Unfortunately, the QFER fit was statistically insignificant (Appendix V), and thus the results from the QFER-φ analysis should be regarded as a preliminary estimate of the φ values. During the development stage of an analytical method, attempting positive interpretations of existing data is crucial to demonstrate the validity and limitations of the method. In the future, statistically rigorous treatment will be necessary to obtain accurate ϕ values for a specific target system.

We compared the original mutational Φ-value analysis developed by Alan Fersht[38,46,47] and our QFER-φ analysis. The original analysis uses the combination of thermodynamic and kinetic experiments of a single-point mutant to provide information on the local structure formation around the mutated side chain in the TSE. In contrast, the QFER-φ analysis uses the data derived from the backbone amide NMR signals and thus provides information about the main chain. The original ϕ-value analysis has limitations due to its assumptions: no changes in folding paths and the absence of U-state perturbations by a mutation[48,49], whereas the QFER-φ analysis does not require these assumptions since no mutants are necessary. From a practical perspective, one must generate one or multiple amino acid mutations per residue and repeat experiments for many residue positions in the mutational Φ-value analysis, but only one amino acid sequence and one set of experiments suffice in the QFER-φ analysis. From an applied standpoint, experimentally derived QFER-φ values are useful for protein folding simulations[50]. Due to a common framework based on a single parabolic curve, the QFER-φ analysis is expected to generate mutually consistent ϕ values. This feature offers a great advantage for molecular simulation studies. The ϕ-ϕ plot is another experimental application that benefits from this feature[51]. A different type of ϕ-value analysis was proposed, using the NMR relaxation dispersion method[52,53]. The NMR method should be useful, but it requires amino acid mutations and thus the same assumptions as the original mutational Φ-value analysis. The ?-value analysis is another mutational method to probe the transition state[54]. A pair of histidine residues (biHis) is introduced into a protein molecule to create a metal ion binding site, and the stabilizing effects caused by increasing concentrations of metal ions are analyzed. The ψ-value analysis has technical advantages over the mutational Φ-value analysis, but the incorporation of the biHis probes has the risk of changing the folding paths.

In sum, the QFER-φ method covers the shortcomings of the mutational ϕ-value and ψ-value analyses. The greatest advantage of the QFER-φ analysis is that mutated proteins are not needed. Bear in mind that the QFER-φ differs from ϕ_whole_, in the analysis of the general conformational changes (Eq. 58). Another discrepancy is attributable to the difference in the time scales of the observation methods: EXSY NMR, relaxation NMR, and HX experiments were the observation methods used in the 18 REFER plots[35], whereas stopped-flow measurements of tryptophan fluorescence were used in the mutational Φ analysis[55]. Finally, the downside of the QFER-φ method is the requirement for highly accurate per-residue measurements; typically, the error range of individual log *K* (or log *k*) values should be sufficiently small compared to the spread of all log *K* (or log *k*) values.

The insights into the dynamic nature of the conformational changes of protein molecules obtained from the residue-specific thermodynamic and kinetic analyses are summarized in Fig. 10. Conventionally, assuming perfect cooperativity among residues, the two-state exchange of a protein is described by a single set of *K, k*, and *k’* values (Fig. 10A). The transition state of the two-state exchange is uniform and characterized by a single ϕ value. By contrast, the residue-based LFER/QFER provides another view (Fig. 10B). Protein molecules are divided into independent structural units, which are conveniently referred to as ‘foldons’. To be exact, the term ‘foldon’ has been used to refer to a structural unit assembled in a stepwise manner during a folding process[10,11], whereas the meaning of ‘foldon’ used here is a structural unit in the TSE identified in the REFER plots and lacks temporal aspects such as the order of formation. The operational definitions are different, but we use the term foldon as a substitute for ‘foldon-like unit’ in this study for convenience. Each foldon is characterized by a specific set of *K, k*, and *k’* values. This situation represents a mosaic of different degrees of cooperativity among residues[30]. The foldons may have different ϕ values describing the polarization of the TSE; i.e., various degrees of the structure formation of foldons. What is important is that all foldons are *consistently* coupled to achieve smooth conformational changes and interactions, judging from the fact that the residues are on the same line/curve in the REFER plots. Exceptionally, although some residues are located apart from the cluster to which they belong, they are still on the same line/curve (Fig. 8), indicating that they are still consistent and do not disturb smooth folding. Such residues may play special roles in the transition state. Finally, we would like to mention the reanalysis of the HX study of apomyoglobin folding intermediates, which was previously described in detail[35]. The majority of the amino acid residues were on a straight line in the REFER plot, but seven residues deviated from the line, suggesting that these outlier residues are *not* consistent with the other residues due to the formation of a third state/structure. Interestingly, these outlier residues precisely corresponded to the transient translocation of an α-helix in the apomyoglobin folding intermediates, which were first discovered a decade-and-a-half ago by other methods. Taken together, the exceptional residues and outlier residues identified in REFER plots provide useful clues about the TSE of the protein structural changes. Accordingly, they could serve as hot spots for manipulating the dynamic properties of protein molecules.

**Figure 10.**
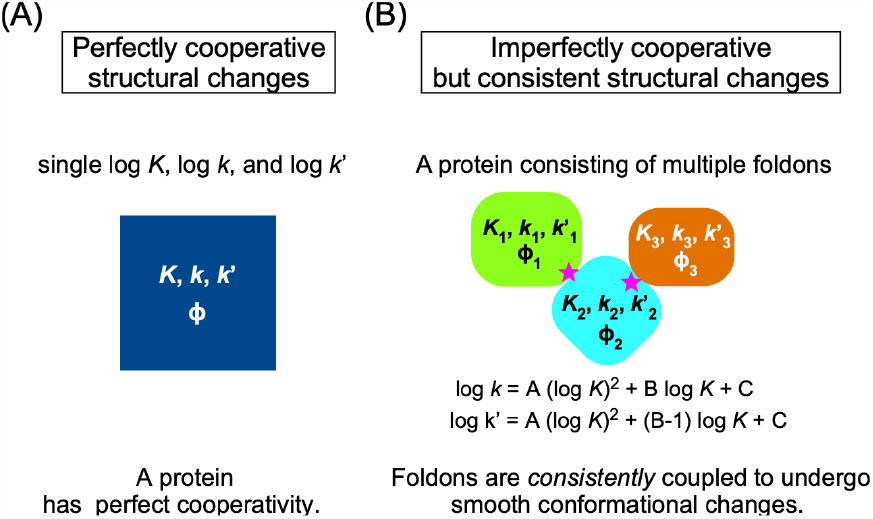
Two views on the dynamic nature of proteins. (A) Conventionally, a protein is assumed to have perfect cooperativity. The two-state exchanges are described with single thermodynamic and kinetic parameters. (B) Residue-based LFER/QFER provides a different view. A protein consists of multiple foldons, which are *consistently* coupled to each other. The magenta stars mark the exceptional residues, which are on the lines/curve in the REFER plot but away from the clusters to which to belong (Fig. 8). They are located at the interfaces between foldons and play special roles in the transition state.

We derived rbLFER and rbQFER from the consistency principle *via* the basic equation (Eq. 37). The Gō model, which embodies the concept of the consistency principle, is an extremely simplified protein model. The three-dimensional protein structure is represented as a chain of points arranged in a two-or three-dimensional lattice, in which only native-centric interactions are effective. Because the lattice model does not contain flexible parts such as loops or tails, the Gō model focuses only on the native contact formation in the core structure. A more realistic coarse-grained model may contain flexible parts, but their behaviors are out-of-focus because non-bonded interactions involving the flexible parts are not included during molecular dynamics simulations. In the field of physics, it is common to simplify or abstract the objects of consideration. This is the reason for the successful derivation of the simple free energy correlations, rbLFER/QFER, in this study. However, when creating REFER plots using experimental data obtained from real protein molecules, the chain entropy of flexible parts, the electrostatic interactions, and the dehydration of hydrophobic residues cannot be ignored.

We consider amino acid residues located in loops and tails. Near the tip of the tail or the center of the loop, the difference in the local environment between the two ground states is small. Therefore, the change in the signal is not large enough to determine the log *K* and log *k* values by using any measurement method (NMR, MS, IR, etc.). The residues in the loops or tails are thus not subject to two-state exchange analysis, so they do not appear as data points in the REFER plot. This consideration explains why rbLFER/QFER, which was derived without consideration of the chain entropy, is compatible with the REFER plots of real proteins possessing flexible parts. We explain this with a specific example, Nukacin ISK-1 (Fig. 8). Except for K1, which provides no cross peak due to fast amide proton exchange with the solvent, the K2, K3, and S4 residues on the tip of the N-terminal tail gave cross peaks in the HSQC NMR spectrum, but the quite similar chemical shifts in the two states prevented the generation of exchange cross peaks in the EXSY NMR spectra. Therefore, the data points in the N-tail portion, K1-K2-K3-S4, do not appear in the REFER plot. For amino acid residues located at the boundary between a tail and a core structure, or at the boundary between a loop and a core structure, the (albeit small) signal changes sense the differences between the two ground states. In some cases, it is possible to determine the log *K* and log *k* values. In the case of nukacin ISK-1, the G5, V6, and I7 residues (P8 is unobservable because proline has no amide hydrogen) at the boundary between the N-tail and the core structure form a cluster in the REFER plot and share similar log *K* and log *k* values. This suggests that the boundary effect is not large enough to disturb the LFER/QFER relationship.

When considering the REFER plot for the core structure, it is also necessary to evaluate the effects of various interactions within the core structure, including the electrostatic interactions and the dehydration of hydrophobic residues. The consistency principle drew the theorem that “*if per-residue log K or log k values have dispersion*, then residue-based LFER/QFER is observable in the REFER plot”. To explain the dispersion of data points in the REFER plot, it is reasonable to consider the influences of electrostatic interactions, the dehydration of hydrophobic residues, and other interactions inside the cores of real proteins as the mechanism for the generation of the per-residue dispersions of log *K* and log *k* values.

A critical challenge remains in demonstrating protein folding examples analyzed with rbLFER/rbQFER. Presently, there are no suitable experimental techniques to obtain accurate residue-level kinetic information on highly skewed equilibrium processes, such as protein folding. We hope that the potential of rbLFER/rbQFER analysis (Fig. 2) will facilitate the development of novel experimental techniques that enable the accurate determination of residue-specific thermodynamic and kinetic parameters for studying the physicochemical basis of protein folding. The combination of ^13^C isotopically edited IR spectroscopy and laser-induced T-jump is one of the promising methods[40,56]

## Appendices Appendix I

The OLS fitting to a quadratic curve, y = Ax^2^ + Bx + C, was performed by the standard polynomial fitting in Excel. The TLS fitting was performed as follows: Let a data point be (x_0_, y_0_) and the foot of the perpendicular be (x, y) from the data point on a quadratic curve, y = Ax^2^ + Bx + C (Fig. 4). When a vector, **u** = (x-x_0_, y-y_0_), and a tangent vector, **v** = (1, 2Ax+B), bisect each other at right angles, then the inner product, **u**·**v** = (x - x_0_) + (y - y_0_)(2Ax + B) is equal to 0, which is a cubic equation in terms of x with real coefficients. The foot point (x, y) was obtained by solving the cubic equation with Cardano’s method. Then, we used the Solver in Excel to find optimal A, B, and C values by minimizing the sum of the square of the orthogonal distance, |**u**|.

## Appendix II

Let *n* be the number of observed residues. The combinatorial number of two residues, *N*, is equal to ½*n*(*n*-1). First, we compute the average of ρ_*ij*_.

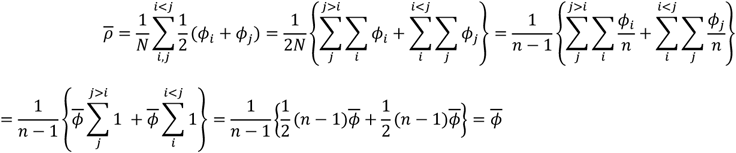

Then, we compute the variance of ρ_*ij*_.

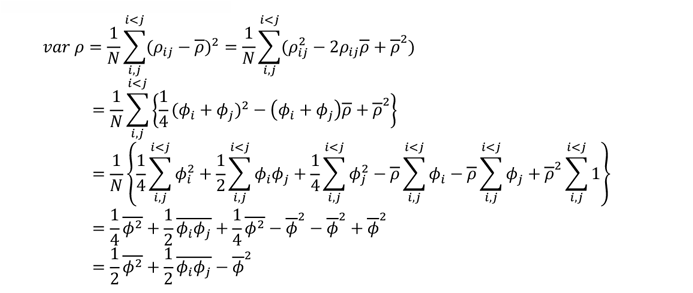

Note that we used the following equalities.

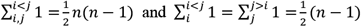

## Appendix III

We solve the next differential equation,

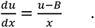

Variable separation gives

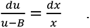

Integrate both sides

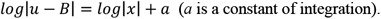

By setting A = ±e^a^, then we have

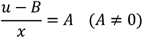

Here, if A is assumed to be 0, then y = Bx is the solution of the differential equation. Thus, A is an arbitrary constant. Using u = y/x = {*f*(x) – C}/x, we obtain the following.

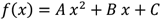

## Appendix IV

The energy barrier height of the slow two-state exchange of nukacin ISK-1 was originally reported to be 10-12*k*_*B*_T[30], but later corrected to 16*k*_*B*_T[44]. However, we found that the original value, 10*k*_*B*_T, is correct. This is because we misinterpreted the empirical value (4 × 10^4^ s^-1^) of the pre-exponential factor of the Arrhenius equation as the *κk*_B_/*h* value of the pre-exponential factor, *κk*_B_*T*/*h*, of the Eyring equation[57].

## Appendix V

*F* is 1.14 < *F*_1,11_=4.84 when the significance level α is set to 0.05. Thus, the QFER fit (Fig. 6) is not statistically significant at α = 0.05. If we relax the significance level α to reject the null hypothesis “QFER is not valid”, *F* is 1.14 ≈ *F*_1,11_=1.13 (α = 0.31).

## Appendix VI

The assignment of log *K* = log *k* – log *k’* to log *k* = A(log *K*)^2^ + B log *K* + C and the simplification produces a quadratic formula, A(log *k*)^2^ - 2A(log *k*)(log *k’*) + A(log *k’*)^2^ + (B-1)log *k* - Blog *k’* + C = 0. This is a conic curve equation, ax^2^ + bxy + cy^2^ + dx + ey + f = 0. The discriminant of the equation is D = b^2^ - 4ac. In our case, D = (−2A)^2^ - 4AA = 0, and thus the quadratic formula represents a tilted parabola.

## Appendix VII

The slope of the curve defined by Eq. 56 is (log *K*^‡^)’ = (2A/ln10) / {0.5(1-0.5)} = 3.47A at ϕ = 0.5. This equation provides the slope of 2.0 for nukacin ISK-1 at ϕ = 0.5. If we apply linear approximation in the range of 0.1 ≤ ϕ ≤ 0.9, then the slope μ is 2.5. The μ value is 25% larger than the slope of the curve at ϕ = 0.5. Thus, we can use an empirical value, μ ≈ 4A.

## Author Contributions

Conceptualization and Investigation, D.K., H.S., and D.F.; Writing – Original Draft, D.K.; Writing – Review & Editing, D.K., H.S., and D.F.; Funding Acquisition, D.K.

### Preprint Server

A preliminary version of this work was deposited in bioRxiv (doi: https://doi.org/10.1101/2022.10.24.513469) on August 21, 2023.

## Acknowledgments

This work was partly performed in the Medical Research Center Initiative for High Depth Omics, at the Medical Institute of Bioregulation, Kyushu University. This work was supported by the Japan Society for the Promotion of Science (JSPS, Japan), KAKENHI Grant Number JP21H02448, and by the Mitsubishi Foundation (Japan) Research Grants in the Natural Sciences, Grant Number 202110017, to D.K.

---

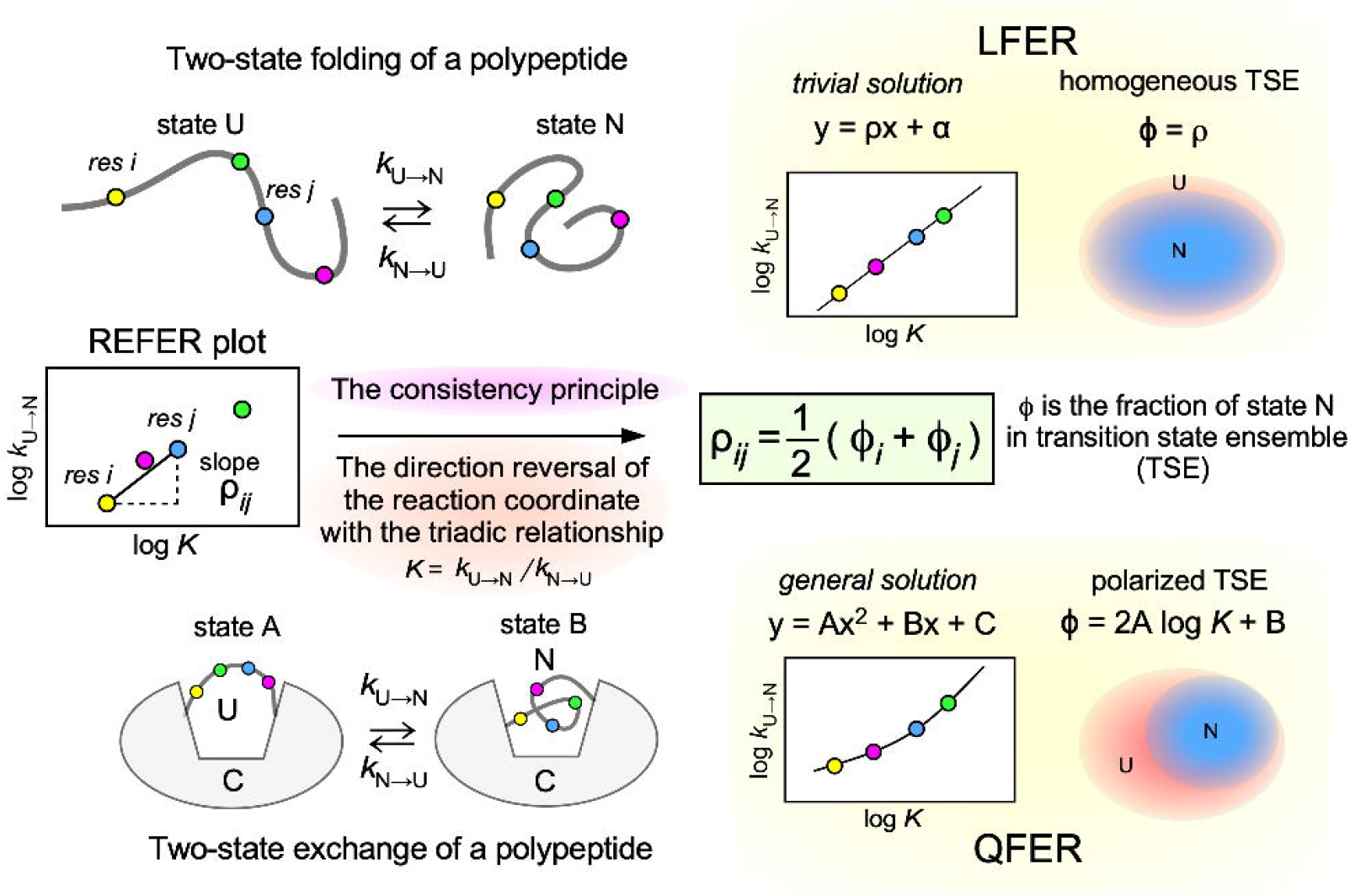

